# Carbon and nitrogen stable isotopic profiling of chimpanzees and other primate hairs in Kalinzu Forest Reserve, Uganda

**DOI:** 10.1101/2025.06.15.658141

**Authors:** Takumi Tsutaya, Natsumi Aruga, Naoto F. Ishikawa, Yoko Sasaki, Haruka Kitayama, Minoru Yoneda, Nana O. Ogawa, Naohiko Ohkouchi, Chie Hashimoto

## Abstract

Stable isotope analysis is a widely used tool in primate ecology for investigating diet and environment, with numerous studies focusing on chimpanzees. However, few studies have explored the dietary niche of chimpanzees in comparison to other primates or examined intra-individual dietary variability. This limitation hinders the understanding of the comparability of isotopic data with the wealth of behavioral observational data in primate ecology. In this study, we report the carbon and nitrogen stable isotope ratios of hairs from wild chimpanzees and four other primate species in the Kalinzu Forest Reserve, Uganda. Bulk analysis revealed that both plant foods and chimpanzees in Kalinzu exhibited lower carbon isotope ratios than expected for the region’s rainfall. Additionally, preliminary compound-specific nitrogen isotope analysis of amino acids was conducted, revealing that chimpanzees in Kalinzu have a lower degree of faunivory than the sympatric *Cercopithecus* primates. Furthermore, ultra-fine sectioning of a hair sample was conducted to investigate dietary variation over daily to weekly timescales. In one adult male chimpanzee, carbon and nitrogen isotope ratios fluctuated by more than 1‰ within approximately 10 days. These findings highlight the importance of recognizing uncontrolled ecological variability and hidden intra-individual dietary changes when interpreting stable isotope data in relation to behavior and environmental traits.

## Introduction

Stable isotopes are used in primate ecological studies to investigate the diet and living environment of various primate species (Crowley et al., 2012; Sandberg et al., 2012). Among them, chimpanzees are probably the most well-studied species, and various aspects of their behaviors and ecology have been revealed mostly by carbon and nitrogen stable isotope analyses (Bădescu et al., 2017; 2022,2025; Fahy et al., 2013; Loudon et al., 2016; Macho and Lee-Thorp, 2014; Oelze et al., 2020; Schoeninger et al., 1999, 2016; Sponheimer et al., 2006; van Casteren et al., 2018). Because of the wealth of behavioral observational evidence, chimpanzees can serve as the model species to evaluate the precision of the stable isotopic methods by comparing the isotopic results with the known behavioral and environmental evidence.

Even in well-studied chimpanzees, however, the scarcity of isotopic research on the two aspects of their diet hinders the accurate interpretations of their isotope ratios as a behavioral and ecological proxy. One aspect is the chimpanzee’s dietary niche compared with other primate species in an ecosystem. Although their diet consists mainly of plants, the contribution of animal foods (faunivory), such as the consumption of insects and vertebrate meat, has rarely been compared with other primate species in the same ecosystem (but see Cerling et al., 2004). While behavioral observation provides a detailed estimate of the time spent feeding on certain food items, stable isotopes reveal the nutritional contribution of some food categories quantitatively, which is an ideal trait for inter-species comparisons (Oelze et al., 2014; Schoeninger et al., 1998). The other understudied aspect is intra-individual dietary variations. Although behavioral observations provide dietary data at a daily resolution, conventional stable isotope analysis only provides an average datum of months or longer periods, generating a gap in evidence between behaviors and isotopes. By investigating how much variation in daily diet affects stable isotope ratios, it would be possible to expand the range of applications for stable isotope analysis in primate ecology.

In this study, profiling on carbon and nitrogen stable isotope ratios (δ^13^C and δ^15^N values, respectively) is performed on wild chimpanzees from Kalinzu Forest Reserve, Uganda, supplemented by that of food plants and other primate species. No previous study has reported stable isotopic data of chimpanzees in Kalinzu, and this study expands the understanding of the ecological determinants of stable isotope ratios of chimpanzees from various sites (Loudon et al., 2016; Schoeninger et al., 2016). The contribution of animal foods is compared between chimpanzees and other sympatric primate species, and compound-specific nitrogen stable isotope analysis was preliminarily applied to further investigate faunivory. Because the trophic enrichment of ^15^N is more pronounced in non-essential amino acids, the δ^15^N values of individual amino acids can be used to obtain more accurate estimates of faunivory (Chikaraishi et al., 2007, 2011; Ohkouchi et al., 2017). Daily to weekly dietary change is also investigated by applying ultra-fine time-series analysis of a chimpanzee’s hair, using an ultra-sensitive analyzer (Ogawa et al., 2010). This profiling deepens understanding of the isotopic ecology of wild chimpanzees and facilitates the connection between behavioral and isotopic proxies.

## Materials and Methods

### Study site

Primate hair and food samples were collected from July 2013 to May 2017 in a moist, medium-altitude evergreen forest in the Kalinzu Forest Reserve, western Uganda, covering an area of 137 km^2^ (30°07′ E, 0°17′ S; altitude 1000–1500 m above sea level) (Hashimoto et al. 1999). The annual rainfall from June 1997 to May 1998 and in 2015 was 1584 mm (Hashimoto et al., 1999) and 1370 mm (Matsuda et al., 2020), respectively. The mean minimum and maximum daily temperatures in each month from November 2013 to April 2016 were 14.0 ± 1.9°C and 27.2 ± 2.1°C, respectively (Matsuda et al., 2020). Two mild rainy seasons (late March–May and late September–December) and two mild dry seasons (January– early March and June–early September) exist. The four major types of vegetation are seen in Kalinzu: mixed mature forest, *Parinari*-dominated mature forest, *Parinari*-dominated secondary forest, and *Musanga*-dominated secondary forest (Hashimoto et al., 1999). Fruits of *Ficus* are abundant throughout the year, providing abundant energy sources to primate species (Furuichi et al., 2001; Hashimoto et al., 2001).

The Kalinzu forest harbors 6 species of diurnal primates: eastern chimpanzee (*P. t. schweinfurthii*), blue monkey (*Cercopithecus mitis*), red-tailed monkey (*Cercopithecus ascanius*), L’Hoest’s monkey (*Allochrocebus.lhoesti*), black and white colobus or guereza (*Colobus guereza*), and the baboon (*Papio anubis*). Primates in Kalinzu have been studied since 1992 (Hashimoto, 1993), and numerous behavioral and ecological studies have been conducted on primate feeding ecology (e.g., Aruga et al., 2015; Furuichi, 2006; Hashimoto et al., 2001; Ihobe, 2001; Matsuda et al., 2020; Tashiro, 2006). However, stable isotopic studies on the diet of primate species in Kalinzu have never been reported.

### Sample collection

Hairs of primates were collected in Kalinzu opportunistically and non-invasively during the study period (Supplementary Table S1). Hairs of chimpanzees were mostly collected from the ground nest or the ground site where grooming was conducted. Hairs of other primate species were mostly collected from dead bodies found inside the forest. The cause of their death seemed to be physical trauma (e.g., falling from a tree or infanticide) in most cases based on the appearance of the bodies and surrounding situations. Chimpanzee hair used for ultra-fine time-series analysis was plucked and collected from the fresh dead body of an adult male (Kobo) who was probably killed by lethal aggression to control the time gap existing in the shed hairs in the telogen phase. Since each hair enters into the telogen phase, where hair incremental growth stops for more than a few months, before shedding (Harkey, 1993; Saitoh et al., 1970), the use of naturally shed hairs complicates the represented time window of the subject hair segments and correspondence between behavioral and isotopic data of the host individual. After the collection, hair was stored in dried or frozen conditions.

Food plant samples were collected during July and August of 2013, February and March of 2015, and October and November of 2015, and they were assigned to dry, dry, and rainy seasons, respectively (Supplementary Table S2). Plant samples were cut into small pieces of approximately less than 1 cm^3,^ air-dried overnight, and dried with silica gel. No sample used in this study was molded. Vertical stratifications that potentially affect plant stable isotope ratios are defined as follows: “crown”, canopy crown; “understray”, the most basal flora below <1.5 m from the ground that does not have direct solar radiation; “subcanopy”, intermediate range between crown and understray that does not have direct solar radiation but locates above 1.5 m from the ground; and “gap”, flora with direct solar radiation because of the clearance of canopy crown. Investigation into the baseline plant isotope ratios is important to accurately interpret the isotope ratios of primates (Wessling et al., 2019) and also benefits DNA-based studies of diet.

The hair and plant samples were exported and analyzed under the material transfer agreement with the Uganda Wildlife Authority and the National Forest Authority. The primate hair samples were exported with the approval of the Uganda National Council for Science and Technology (NS 184) and the Ministry of Economy, Trade and Industry, Japan (17JP000003/TI), under the Convention on International Trade in Endangered Species (CITES). All research procedures followed the Code of Best Practices for Field Primatology (International Primatological Society, 2014) and were approved by the Ethics Committee for Animal Research of the Graduate University for Advanced Studies (SKD2025AR003).

### Sample processing

Hair samples were washed with Milli-Q water under sonication and dried. Then, lipid-extraction was performed by soaking the hairs in a hexane:dichloromethane (3:2 in volume) solution over 2 hours. After the removal of organic solvent with washing by Milli-Q water, the hair was air-dried and used for stable isotope analyses.

Plant samples were freeze-dried overnight and crushed into a fine powder with a stainless motor and pestle, if the matrix heterogeneity was high. The homogenized samples were used for stable isotope analyses.

### Ultra-fine sectioning

A single hair strand in the growing anagen phase of an adult male chimpanzee (Kobo) was cut into 1 mm sections and used for the time series analysis. The hair was wrapped in aluminum foil, wetted with ethanol to prevent electrostatics, and manually dissected in a 1 mm unit with a pair of scissors. To avoid the accumulation of small errors, the hair was first cut into 5 mm segments, and each segment was cut into five 1 mm pieces. When the 1 mm segment was lost, data were not obtained in the corresponding time period. Based on the previous data in humans (Harkey, 1993; Saitoh et al., 1970), the hair growth rate was assumed to be 1 mm in 3 days (i.e., 1 cm in 1 month). Therefore, for example, the first, second, and third 1 mm segments from the hair root represent the 3-day periods of the last 1–3 days (median 1.5 days), 4–6 days (median 4.5 days), and 7–9 days (median 7.5 days) from the estimated date of the death (July 30, 2013).

The microscopic observations on the hair follicle were performed on the hair strands that were in the same appearance as the sectioned strand and were not used for stable isotope analyses. Keyence VHX-2000 was used. The follicles are transparent white colors with an elongated, straight form, which are the typical features of hairs in the anagen phase (Supplementary Figure S1). Therefore, there was no gap between the assigned time window to the segments and the actual timing of hair growth that reflects dietary isotopic signals at the time.

### Bulk stable isotope analysis

Bulk carbon and nitrogen stable isotope ratios of plant samples were measured using an elemental analyzer–isotope ratio mass spectrometry (Thermo Flash 2000 elemental analyzer, Finnigan ConFlo III interface, and Thermo Delta V mass spectrometer) at the University Museum, University of Tokyo, Japan. Carbon and nitrogen isotope ratios were expressed in δ notation relative to standards, δ^13^C (‰) and δ^15^N (‰), with Vienna Pee Dee Belemnite (VPDB) and atmospheric nitrogen (AIR) as the standard for carbon and nitrogen, respectively. The δ^13^C and δ^15^N values were calibrated against the laboratory working standard (L-alanine: δ^13^C = −19.6 ± 0.2‰; δ^15^N = 8.7 ± 0.2‰) provided by SI Science (Saitama, Japan), whose values were determined by the NBS 19 and the International Atomic Energy Agency (IAEA) Sucrose ANU (calibrated against Pee Dee Belemnite and IAEA N1 and IAEA N2 (calibrated against AIR) international standards, respectively. Based on repeated measurements of the calibration standards, analytical errors were determined to be less than ±0.1‰ for both δ^13^C and δ^15^N.

### Ultra-sensitive bulk stable isotope analysis

Bulk carbon and nitrogen stable isotope ratios (δ^13^C_bulk_ and δ^15^N_bulk_) of hair samples were measured using a modified elemental analyzer/isotope mass spectrometer at the Japan Agency for Marine-Earth Science and Technology (JAMSTEC). Instrumentation consisted of an elemental analyzer (Flash EA 1112, Thermo Finnigan) coupled to an isotope ratio mass spectrometer (Delta plus XP, Thermo Finnigan) through a continuous flow interface (ConFloIII, Thermo Finnigan) with modification specifically made to improve the sensitivity of the analysis (Ogawa et al., 2010). Isotope ratios were calibrated using three laboratory standards: L-tyrosine (BG-T: δ^13^C = −20.83 ± 0.10‰, δ^15^N = 8.74 ± 0.04‰), L-proline (BG-P: δ^13^C = −10.27 ± 0.04‰, δ^15^N = 13.51 ± 0.02‰), and DL-alanine (CERKU-01: δ^13^C = −25.36 ± 0.08‰, δ^15^N = −2.89 ± 0.04‰). These isotope compositions were calibrated with authentic standards (Tayasu et al., 2011). The analytical errors determined by the repeated measurements of L-tyrosine (BG-T) at the same timing of sample analysis were± 0.2‒0.7‰ and ± 0.2‒0.6‰ for δ^13^C and δ^15^N, respectively.

### Compound-specific stable isotope analysis of amino acids

Compound-specific isotope analysis of amino acid δ^15^N (δ^15^N_AA_) was performed for selected hair samples. Samples for CSIA were prepared based on the amino acid derivatization procedures described in Chikaraishi et al. (2015). Briefly, samples (∼1 mg) were hydrolyzed with 12 M HCl at 110 °C overnight, and then washed with *n*-hexane:dichloromethane (3:2 in volume) to remove the remaining hydrophobic compounds. After the hydrolysis, samples were derivatized using thionyl chloride:2-propanol (1:4 in volume) at 110 °C for 2 h and then pivaloyl chloride:dichloromethane (1:4 in volume) at 110 °C for 2 h. Then, the amino acid derivatives were extracted with *n*-hexane:dichloromethane (3:2 in volume). The δ^15^N values of each amino acid were determined using a gas chromatograph (6890 N, Agilent Technologies) coupled to an isotope ratio mass spectrometer (Delta plus XP, Thermo Finnigan) via a GC-combustion III interface (Thermo Finnigan) at JAMSTEC (Ishikawa et al., 2018, 2022). Reference mixtures of six amino acids (alanine, valine, leucine, norleucine, aspartic acid, glutamic acid, and phenylalanine, provided by Indiana University and Shoko Science Co., Ltd.) with known δ^15^N values (ranging from +1.7‰ to +45.7‰) were analyzed every six sample runs. Duplicate or triplicate measurements were performed for each amino acid. The analytical error of the standards was typically smaller than 1.0‰.

### Behavioral observation

Behavioral observations of chimpanzees were performed by local research assistants. The scan sampling of behaviors of chimpanzees in the observer’s sight was obtained every 10 minutes from 8:00 to 17:00 from Monday to Saturday. The species and parts of food items consumed were also recorded. The presence of Kobo, the chimpanzee individual whose hair was used for ultra-fine time-series analysis, and his feeding items were evaluated from this group-scan-based behavioral observation records.

Mostly based on these data, each combination of plant species and part was scored on whether it is consumed by each primate species. The scoring results were reviewed by local research assistants and researchers working on these primate species in Kalinzu. Results reported by Matsuda et al. (2020) were used for food items in guereza.

### Data analysis

Statistical analyses were performed in R software environment, version 4.2.3 (R Core Team, 2023). Linear mixed models (LMMs) were applied using *lme4*, version 1.1-31 (Bates et al., 2015) and *lmerTest*, version 3.1-3 (Kuznetsova et al., 2017) packages. The statistical significance level was set as α < 0.05. Liner mixed models were applied with the following conditions:

- To reveal the drivers of isotopic variations in plants, plant δ^13^C_bulk_ or δ^15^N_bulk_ values were set as the response variable, and part (fruits, leaves, pith, or others), stratification (crown, subcanopy, understray, or gap), season (dry or rainy) were set as the explanatory variables. The combination of species and part was set as random effect.
- To reveal the isotopic difference of the primate species, hair δ^13^C_bulk_ or δ^15^N_bulk_ values were set as the response variable, and primate species and age class (infant or non-infant) were set as response variables. The individual ID of primates was set as a random effect.

To reduce the sampling bias, mean values of samples in the same category (i.e., individuals in hairs and the combination of species and part in plants) were used to calculate summary statistics, such as means and standard deviations, and when applying statistical tests. On the other hand, bias in the number of samples was considered in LMMs by incorporating them into random effects. Also, to control the effect of breast milk consumption to the isotope ratios (Tsutaya and Yoneda, 2015), only non-infant data were used for comparison.

The trophic position (TP) was calculated based on the stable nitrogen isotope ratios of glutamic acid and phenylalanine with the following equation proposed in Chikaraishi et al. (2011, 2014):

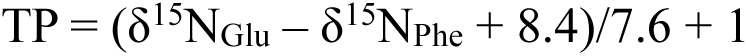

Uncertainty in the TP was calculated by propagation of errors by combining the uncertainties in the isotopic difference between glutamic acid and phenylalanine in reference mixtures of amino acids (σ = 1.5‰), the trophic discrimination factors of glutamic acid and phenylalanine during each trophic transfer (σ = 1.2‰ and 0.5‰, respectively: Chikaraishi et al. 2009), and the measured analytical reproducibility of δ^15^N_Glu_ and δ^15^N_Phe_ values in each sample, by using the equation shown in Ishikawa et al. (2022). It should be noted that the propagated errors might overestimate the TP estimate errors.

## Results

### Bulk stable isotope ratios of plants

The total averages of δ^13^C_bulk_ and δ^15^N_bulk_ values of each combination of plant species and part that were consumed by any of the primate species were -30.1 ± 3.1‰ and 3.5 ± 1.1‰, respectively (Table 1; Figure 1a). The average δ^13^C_bulk_ (from -30.2‰ to -28.8‰) or δ^15^N_bulk_ (from 3.0‰ to 3.8‰) values of consumed food items were similar among each primate species (Table 1).

**Figure 1.**
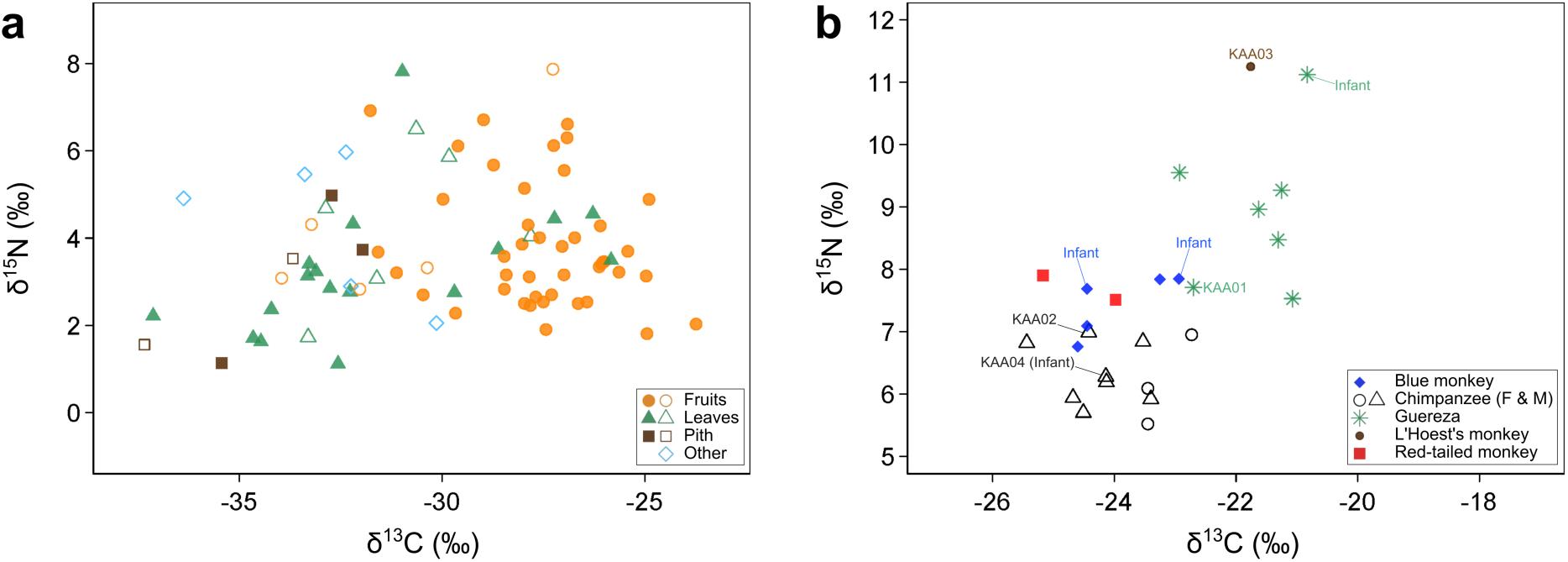
Distributions of carbon and nitrogen stable isotope ratios of **a**) food plants and **b**) primate hairs from Kalinzu. Plant species and part eaten by chimpanzees are shown in solid points (**a**). Samples used for CSIA are shown with its ID (KAA) and those from infants are also shown (**b**).

**Table 1.** Summary of stable isotope ratios of plant foods of the Kalinzu.

| Consumer | Part | %C | | %N | | $\delta^{13}\text{C}$ | | $\delta^{15}\text{N}$ | | n |
| --- | --- | --- | --- | --- | --- | --- | --- | --- | --- | --- |
|  |  | Mean | SD | Mean | SD | Mean | SD | Mean | SD |  |
| Total | Flower | 41.8 | — | 2.7 | — | -31.2 | — | 2.5 | — | 1 |
|  | Fruit | 44.3 | 1.6 | 1.7 | 0.8 | -28.0 | 1.9 | 3.7 | 1.1 | 16 |
|  | Leaves | 44.1 | 3.2 | 3.3 | 1.2 | -31.8 | 2.6 | 3.3 | 1.2 | 12 |
|  | Pith | 40.8 | 4.6 | 2.8 | 1.6 | -34.4 | 1.5 | 2.9 | 0.5 | 2 |
|  | Seed | 44.3 | — | 2.8 | — | -34.0 | — | 5.4 | — | 1 |
| Chimpanzee | Fruit | 44.6 | 2.1 | 1.6 | 0.8 | -27.5 | 1.5 | 3.6 | 1.0 | 14 |
|  | Leaves | 44.8 | 2.5 | 3.2 | 1.3 | -31.7 | 2.9 | 3.2 | 1.1 | 9 |
|  | Pith | 44.0 | — | 1.7 | — | -33.4 | — | 3.3 | — | 1 |
|  | **Total** | **44.6** | **2.2** | **2.2** | **1.2** | **-29.3** | **3.0** | **3.4** | **1.0** | **24** |
| Blue monkey | Fruit | 45.0 | 1.8 | 1.7 | 0.6 | -27.7 | 1.4 | 3.7 | 1.3 | 9 |
|  | Leaves | 43.2 | 4.2 | 2.8 | 1.2 | -30.9 | 1.7 | 3.8 | 0.9 | 5 |
|  | **Total** | **44.4** | **2.9** | **2.1** | **1.0** | **-28.8** | **2.1** | **3.7** | **1.1** | **14** |
| Guereza | Flower | 41.8 | — | 2.7 | — | -31.2 | — | 2.5 | — | 1 |
|  | Fruit | 45.5 | 0.1 | 1.9 | 1.4 | -25.7 | 1.1 | 3.2 | 0.5 | 2 |
|  | Leaves | 44.4 | 2.4 | 3.5 | 1.7 | -31.7 | 3.9 | 3.0 | 1.1 | 5 |
|  | **Total** | **44.3** | **2.2** | **3.0** | **1.5** | **-30.2** | **4.1** | **3.0** | **0.9** | **8** |
| L'Hoest's monkey | Flower | 41.8 | — | 2.7 | — | -31.2 | — | 2.5 | — | 1 |
|  | Fruit | 45.1 | 1.9 | 1.8 | 0.6 | -28.2 | 2.1 | 3.8 | 1.2 | 9 |
|  | Leaves | 42.4 | 5.2 | 3.3 | 1.2 | -31.0 | 1.6 | 4.0 | 1.3 | 3 |
|  | Pith | 40.8 | 4.6 | 2.8 | 1.6 | -34.4 | 1.5 | 2.9 | 0.5 | 2 |
|  | Seed | 44.3 | — | 2.8 | — | -34.0 | — | 5.4 | — | 1 |
|  | **Total** | **43.8** | **3.2** | **2.3** | **1.0** | **-30.0** | **3.0** | **3.8** | **1.2** | **16** |
| Red-tailed monkey | Fruit | 45.0 | 1.8 | 1.7 | 0.6 | -27.7 | 1.4 | 3.7 | 1.3 | 9 |
|  | Leaves | 43.4 | 4.8 | 2.9 | 1.4 | -31.5 | 1.3 | 3.9 | 1.1 | 4 |
|  | **Total** | **44.5** | **2.9** | **2.1** | **1.1** | **-28.8** | **2.2** | **3.7** | **1.2** | **13** |

The results of LMMs showed significantly higher δ^13^C_bulk_ values of fruits compared with leaves, significantly lower δ^13^C_bulk_ values in samples obtained from the subcanopy compared with those from the crown, significantly lower δ^15^N_bulk_ values in piths and other parts compared with fruits, and significantly higher δ^15^N_bulk_ values of samples from the understray compared with those from the crown (Table 2). The average δ^13^C_bulk_ value of fruits was 3.8‰ higher than that of leaves (Table 1). Compared with plant samples obtained from the crown, the average δ^13^C_bulk_ value of plant samples obtained from the subcanopy was 2.5‰ lower, and the average δ^15^N_bulk_ value of plant samples obtained from the understray was 0.3‰ higher (Supplementary Table S3).

**Table 2.**
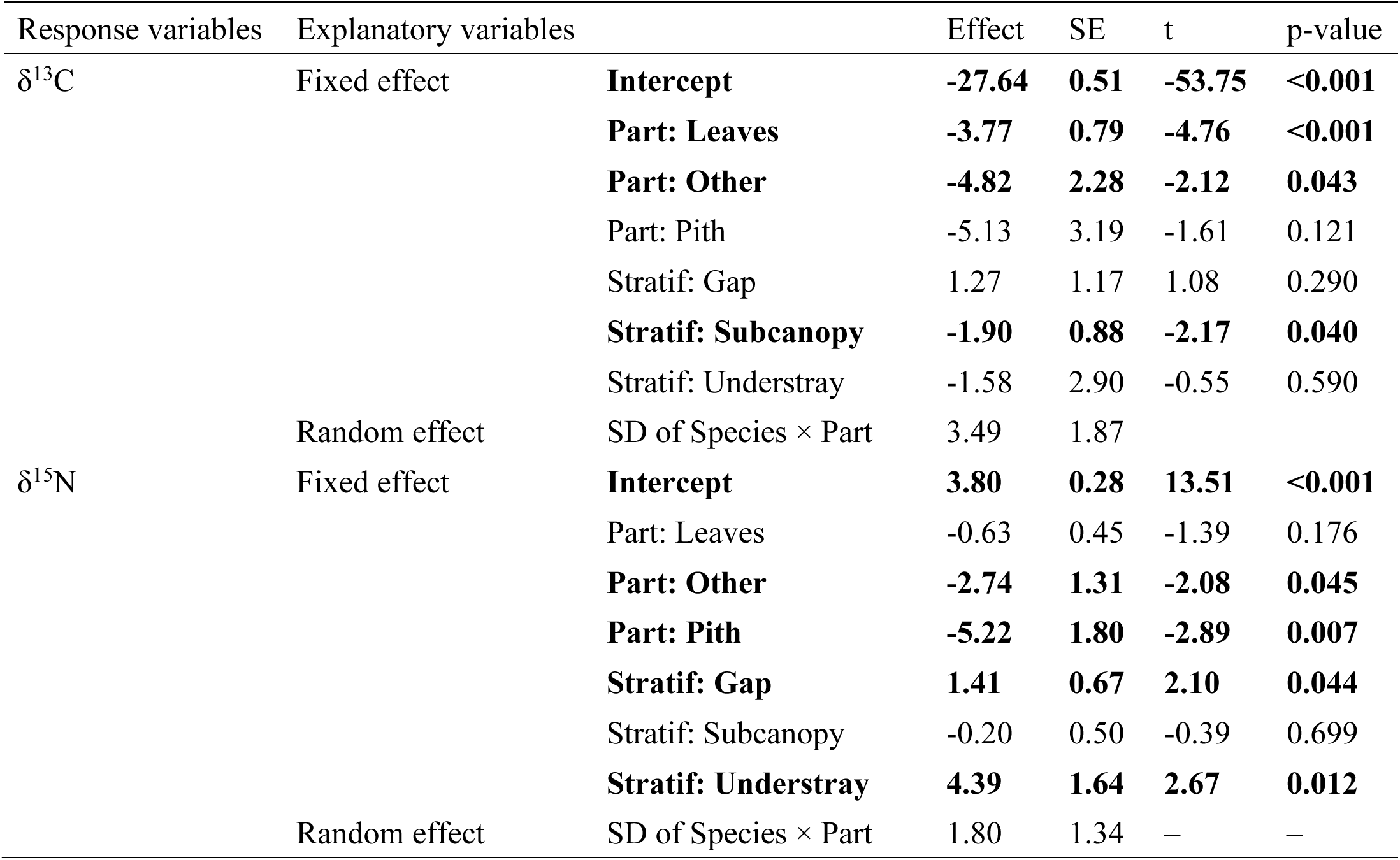
Explanatory variables and intercept in the linear mixed models for plant stable isotope ratios in the Kalinzu.

### Bulk stable isotope ratios of primate hairs

The average bulk stable isotope ratios of five primate species ranged from -24.6‰ to - 20.8‰ in carbon and from 6.2‰ to 11.3‰ in nitrogen (Table 3; Figure 1b). The results of LMMs showed significantly higher δ^13^C_bulk_ values of guereza and L’Hoest’s monkey compared with chimpanzees, significantly higher δ^15^N_bulk_ values of guereza, L’Hoest’s

**Table 3.** Summary of stable isotope ratios of primates hair of the Kalinzu.

| Species | Category | %C | | %N | | $\delta^{13}\text{C}$ | | $\delta^{15}\text{N}$ | | n |
| --- | --- | --- | --- | --- | --- | --- | --- | --- | --- | --- |
|  |  | Mean | SD | Mean | SD | Mean | SD | Mean | SD |  |
| Blue monkey | Infant | 47.8 | 2.1 | 15.9 | 0.4 | -23.7 | 1.1 | 7.8 | 0.1 | 2 |
|  | Non-infant | 45.5 | 3.5 | 15.8 | 1.7 | -24.1 | 0.7 | 7.2 | 0.6 | 3 |
| Chimpanzee | Infant | 40.7 | – | 14.3 | – | -24.1 | – | 6.3 | – | 1 |
|  | Non-infant F | 38.0 | 1.9 | 12.7 | 0.6 | -23.2 | 0.4 | 6.2 | 0.7 | 3 |
|  | Non-infant M | 42.7 | 2.5 | 14.1 | 1.0 | -24.1 | 0.6 | 6.4 | 0.5 | 5 |
|  | Non-infant total | 40.9 | 3.2 | 13.6 | 1.1 | -23.8 | 0.7 | 6.3 | 0.6 | 8 |
| Guereza | Infant | 48.9 | – | 17.3 | – | -20.8 | – | 11.1 | – | 1 |
|  | Non-infant | 41.0 | 2.5 | 14.0 | 0.9 | -21.8 | 0.8 | 8.6 | 0.8 | 6 |
| L'Hoest's monkey | Non-infant | 45.5 | – | 16.0 | – | -21.8 | – | 11.3 | – | 1 |
| Red-tailed monkey | Non-infant | 47.3 | 2.2 | 16.5 | 0.4 | -24.6 | 0.8 | 7.7 | 0.3 | 2 |

monkey, and red-tailed monkey compared with chimpanzees, and significantly higher δ^15^N_bulk_ values of infants (Table 4).

**Table 4.**
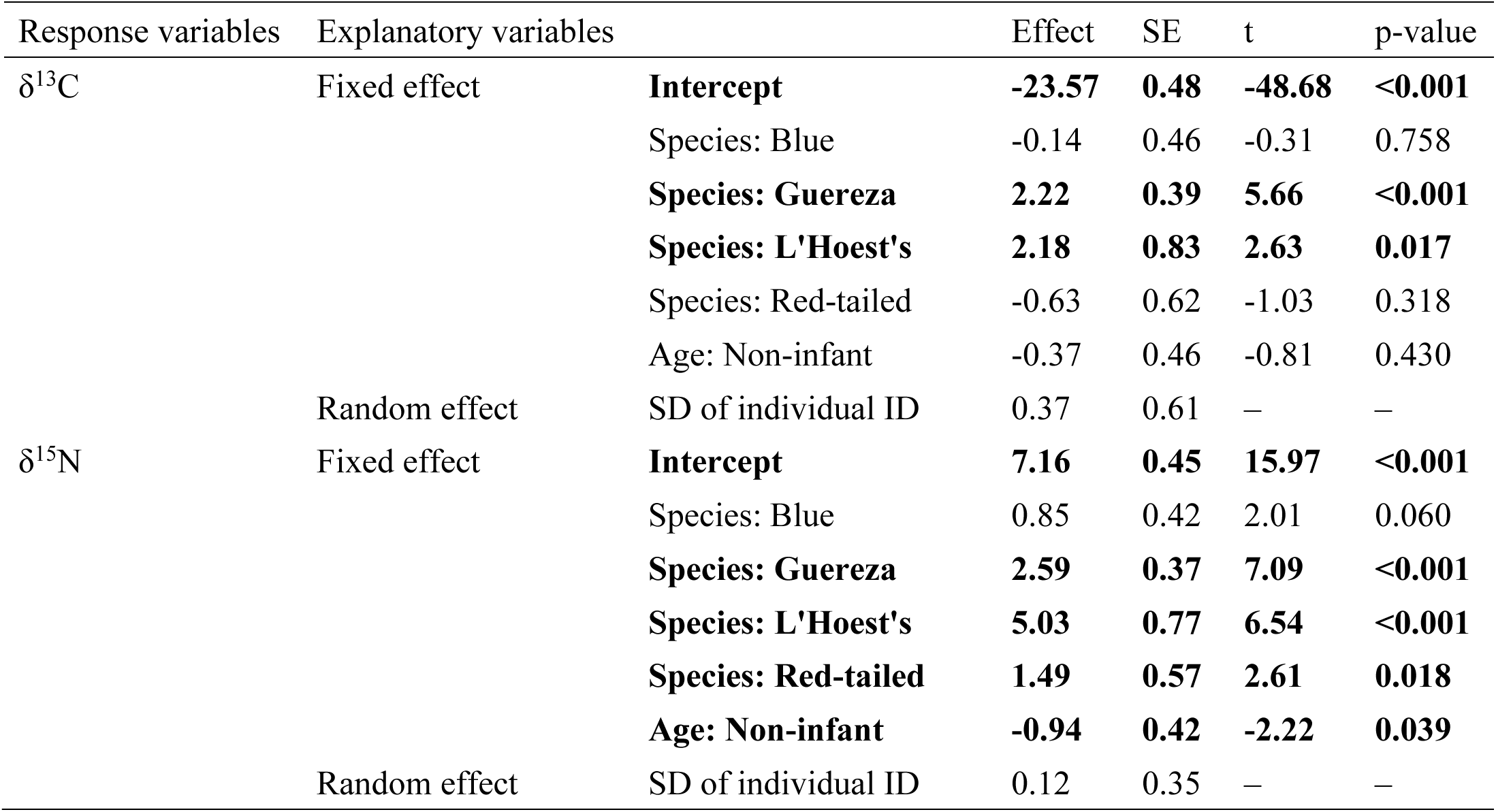
Explanatory variables and intercept in the linear mixed models for primate hair stable isotope ratios in the Kalinzu.

Mann–Whitney U-tests showed no significant sex difference in δ^13^C_bulk_ (U = 13, p = 0.134) and δ^15^N_bulk_ (U = 6, p = 0.786) values of chimpanzees.

The average δ^13^C_bulk_ or δ^15^N_bulk_ values of the non-infant Kalinzu chimpanzees were -23.8 ± 0.7‰ and 6.3 ± 0.6‰, respectively (Table 3). The isotopic differences between diet and hair were 5.5‰ and 2.9‰ for carbon and nitrogen, comparable to those estimated in captive chimpanzees in an experimental setting (Tsutaya et al., 2017). Comparison of these values with the isotope ratios of chimpanzees and their food plants from other study sites across Africa indicated lower δ^13^C_bulk_ values for both hair and food plants but similar δ^15^N_bulk_ values compared with those predicted by the relationships between annual rainfall and isotope ratios (Figure 2; Supplementary Table S4). Although the δ^13^C_bulk_ values of hair and plants were 1.7‰ and 1.3‰ lower than the predicted values, respectively, the δ^15^N_bulk_ values were both only 0.1‰ higher than the predicted values (Figure 2).

**Figure 2.**
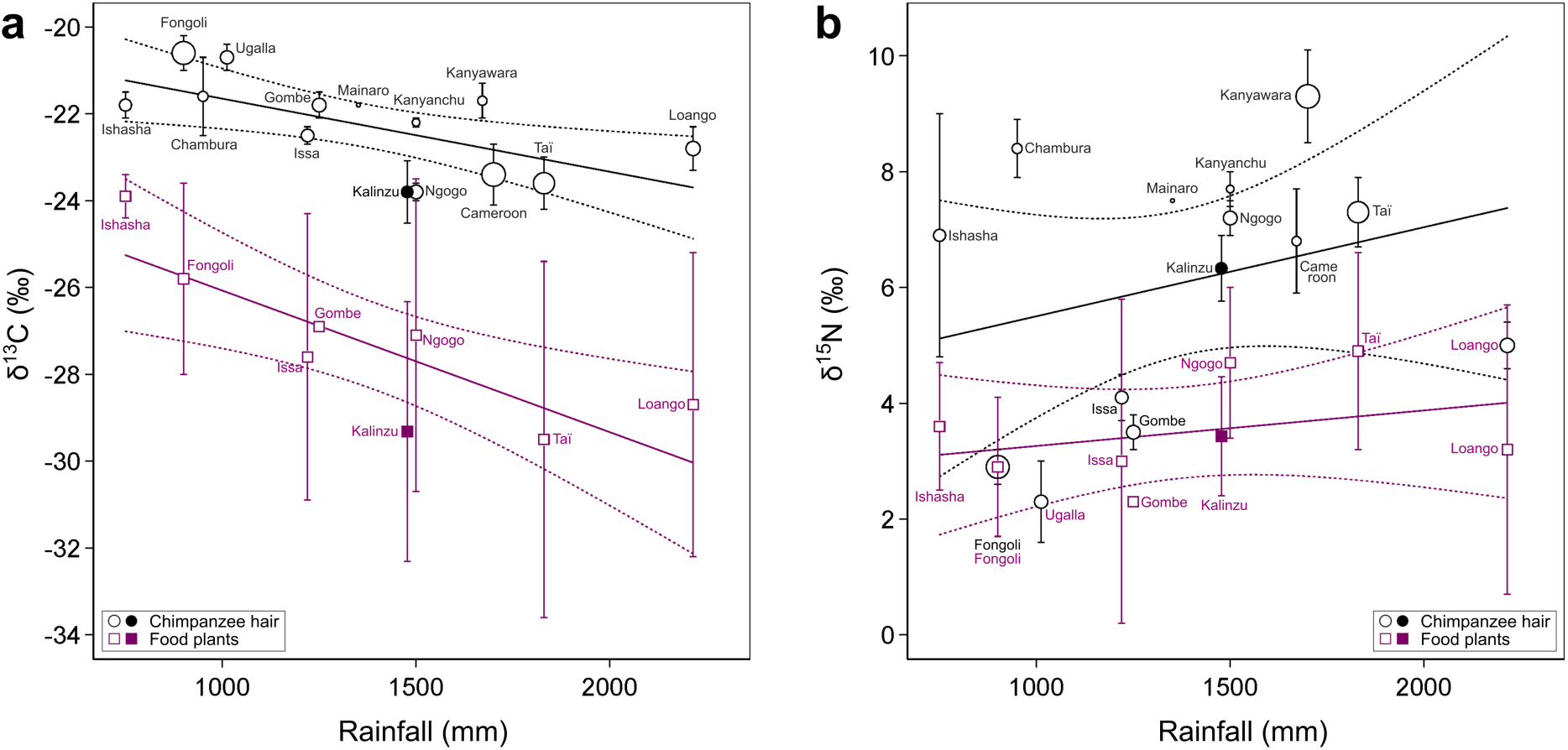
The relationship between mean annual rainfall and **a**) carbon or **b**) nitrogen isotope ratios of food plants and hairs of chimpanzees across different sites. The 1SD ranges of each data point are shown for isotope ratios. The diameters of plots for hairs corresponds to the number of analyzed samples in each study. The predicted relationship and its error ranges are shown with solid and dotted lines, respectively. Please see Supplementary Table S4 for the information on each site and Supplementary Text S1 for the details of the linear regression.

### Time-series analysis of a chimpanzee hair

Ultra-fine time-series analysis of Kobo’s hair revealed unexpected rapid fluctuations of δ^13^C_bulk_ or δ^15^N_bulk_ values in daily to weekly scales (Figure 3; Supplementary Table S5). The δ^13^C_bulk_ or δ^15^N_bulk_ values of 1 mm segments corresponding approximately 3 days during 84 days before the death ranged from -24.7‰ to -23.7‰ and from 6.5‰ to 8.0‰ (Supplementary Table S5). Although most parts of the hair segments have a relatively constant isotope ratio, a 1.0‰ increase occurred during 9 days just before the death in δ^13^C_bulk_ values, and a 1.4‰ increase and subsequent 1.0‰ decrease occurred during 12 days in δ^15^N_bulk_ values (Figure 3). If the isotope ratios of 1 mm segments were averaged, based on the length, into hypothesized 5 mm segments, such rapid fluctuations were not seen (Figure 3), and the difference between the minimum and maximum values decreased to 0.4‰ and 0.6‰ for δ^13^C_bulk_ or δ^15^N_bulk_ values, respectively (Supplementary Table S6).

**Figure 3.**
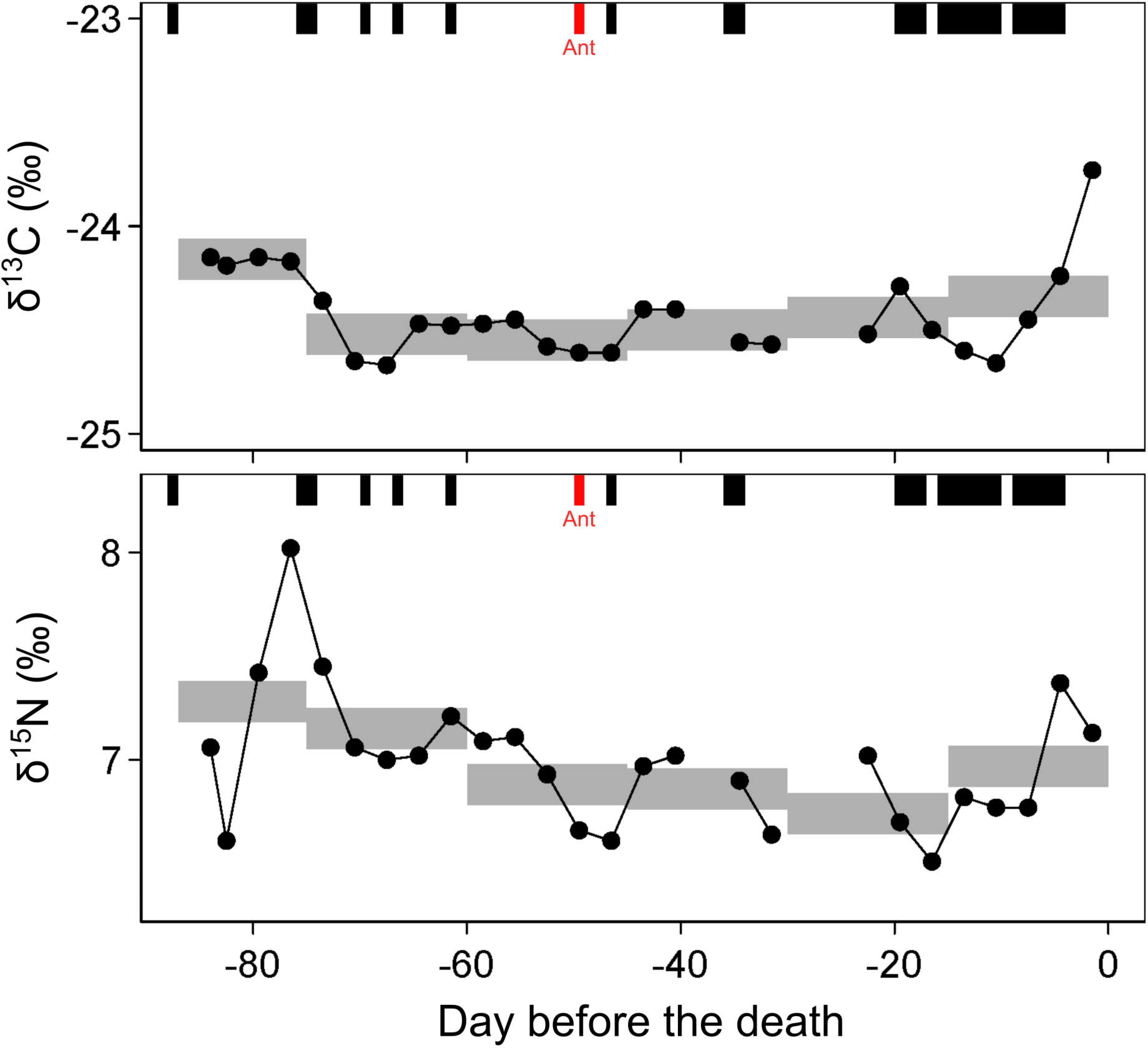
Carbon and nitrogen isotope ratios of the ultra-fine sections of the hair of a male chimpanzee (Kobo), plotted against the calculated days before the death. The average values of hypothesized 5 mm segments are shown in gray horizontal bars with the height of typical analytical uncertainty of ±0.1‰. Days with behavioral observation records are shown in rectangular blocks above the isotopic trajectories, and days with the observation of faunivory are shown in red.

Among the 29 days in which Kobo was observed during the 87 days period before his death, animal food consumption (i.e., ant-eating) was only seen once at 49 days before his death, and no rapid δ^13^C_bulk_ or δ^15^N_bulk_ fluctuation can be seen after this ant-eating event. Kobo was relatively well observed during the 19-day period before his death, in which the rapid increase in hair δ^13^C_bulk_ values occurred, but no animal food consumption was recorded.

### Compound-specific isotope analysis of primate hairs

The measured δ^15^N_Phe_ values of chimpanzees, guereza, and L’Hoest’s monkey were similar (from 12.3‰ to 13.4‰), indicating that they share the same nitrogen source (Figure 4; Table 6; Supplementary Table S7). It should be noted that phenylalanine for guereza and L’Hoest’s monkey coeluted with unknown impurities in one of their three chromatograms. Consequently, their δ^15^N_Phe_ values were determined in duplicate, and guereza δ^15^N_Phe_ particularly showed larger uncertainty than others (Figure 4). The δ^15^N_Glu_ values, which typically increase with the increase in TP, were the lowest in chimpanzees (8.8‰ and 9.7‰) and larger in guereza (12.5‰) and in L’Hoest’s monkey (15.3‰) (Table 5).

**Figure 4.**
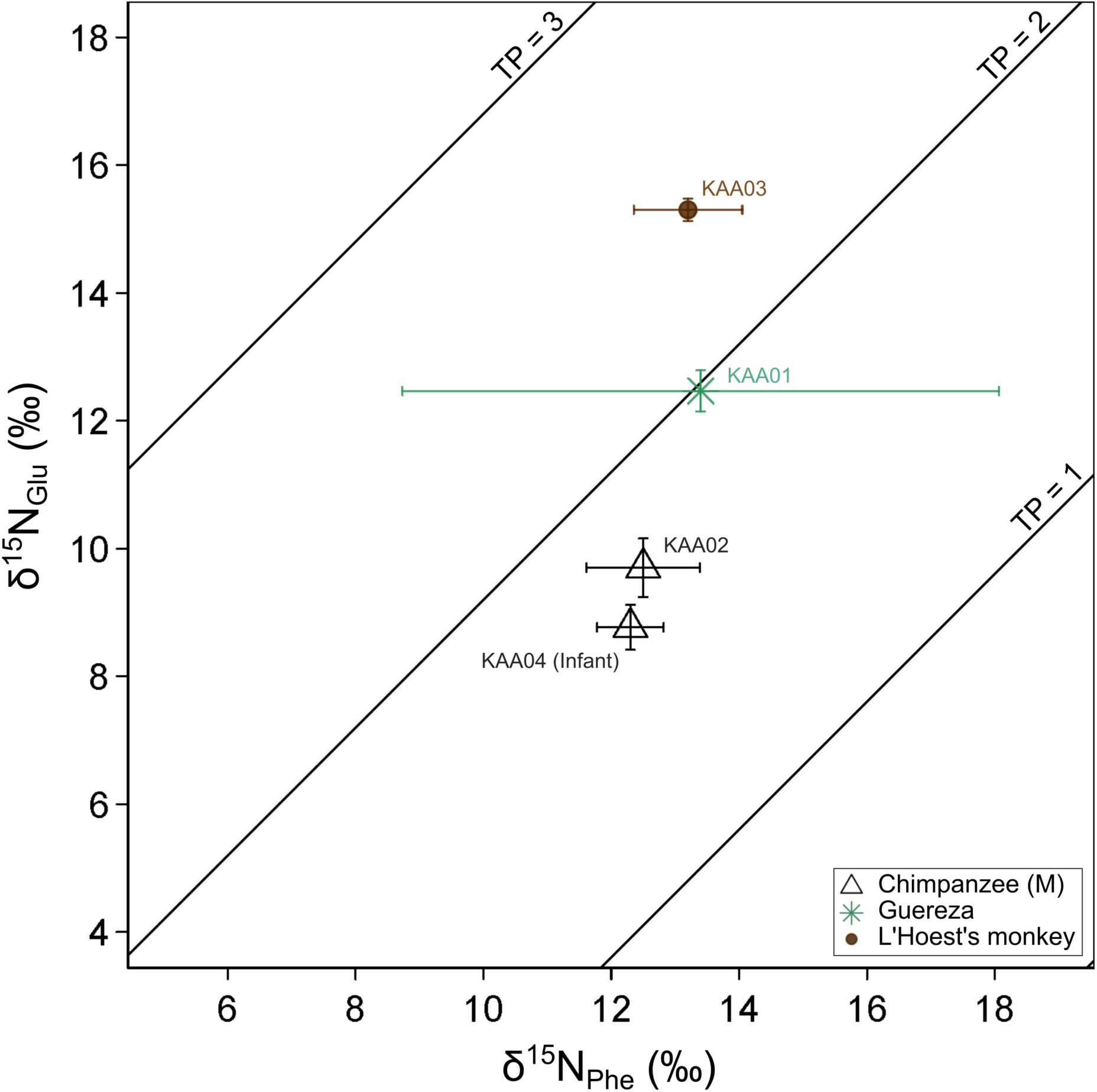
Nitrogen stable isotope ratios of Glutamic acid and Phenylalanine of primate hairs from Kalinzu. Sample IDs and estimated TPs calculated based on the equation mentioned in the main text are also shown

**Table 5.** Summary of the stable isotope ratios of amino acids of primate hair samples from the Kalinzu Forest Reserve.

| Species | ID | $\delta^{15}\text{N}_{\text{Glu}}$ | | $\delta^{15}\text{N}_{\text{Phe}}$ | | TP | |
| --- | --- | --- | --- | --- | --- | --- | --- |
|  |  | Mean | SD | Mean | SD | Mean | Propagated error |
| Chimpanzee | KAA02 | 9.7 | 0.5 | 12.5 | 0.9 | 1.7 | 0.27 |
|  | KAA04 | 8.8 | 0.4 | 12.3 | 0.5 | 1.6 | 0.24 |
| Guereza | KAA01 | 12.5 | 0.3 | 13.4 | 4.7 | 2.0 | 0.67 |
| L’Hoest’s monkey | KAA03 | 15.3 | 0.2 | 13.2 | 0.9 | 2.4 | 0.33 |

The TPs calculated from the results of CSIA were 1.6–1.7, 2.0, and 2.4 for chimpanzees, guereza, and L’Hoest’s monkeys, respectively (Table 5; Supplementary Figure S2). The TPs are expected to be 2.0 for obligate plant consumers and 3.0 for faunivorous animals that exclusively consume plant-eating animals. This preliminary result indicates that the major dietary nitrogen sources were plants in chimpanzees and guereza, but certain amounts of foods in higher TP, such as insects, were incorporated into the diet of L’Hoest’s monkey, although more measurement data are needed to reach a robust conclusion.

## Discussion

Factors affecting stable isotope ratios are investigated at various levels in Discussion. We do not employ hypothesis testing here, but exploratory data investigation can illuminate important areas for further study. Such detailed knowledge of primate diet also contributes to a better interpretation of results obtained from DNA-based studies, such as fecal metagenomics (Mallot et al., 2017) and chemosensory gene evolution (Hayakawa et al., 2012; Itoigawa et al., 2021), by providing quantitative baseline data of their feeding ecology.

### Isotopic variation among ecosystems

The hair and plant food stable isotope ratios of Kalinzu chimpanzees were generally consistent with those predicted based on the relationship between rainfall and isotope ratios, although the hair and plant δ^13^C_bulk_ values were lower than those predicted from the general trend across various chimpanzee study sites (Figure 2; Supplementary Table S4). The δ^13^C_bulk_ values of terrestrial C3 plants show a strong negative correlation with the amount of rainfall (Kohn, 2010), and chimpanzee hairs also reflect this trend (Loudon et al., 2016; Schoeninger et al., 2016). Because both the hair and plant of chimpanzees from Kalinzu show lower δ^13^C_bulk_ values, the deviation from the general trend was not caused by the distinct dietary habits of Kalinzu chimpanzees but by environmental factors affecting the baseline plant isotope ratios. Although the cause of this baseline change is unknown, this result emphasizes the importance of analyzing the isotope ratios of food plants to compare those of primate tissues (Wessling et al., 2019).

Multiple factors affect plant δ^15^N_bulk_ values, such as climate, type of mycorrhizal association, and nitrogen sources (Amundson et al., 2003; Craine et al., 2009, 2015; Szpak, 2014), and the relationship between plant δ^15^N_bulk_ values and environmental factors are complex. Reflecting such complexity, chimpanzee δ^15^N_bulk_ values largely vary by baseline change and dietary content, such as meat (Loudon et al., 2016; Schoeninger et al., 2016). Despite this large variation, plant and chimpanzee δ^15^N_bulk_ values did not seem to deviate from typical values previously reported, suggesting that animal meat, typically showing higher δ^15^N values than plants, was not the major dietary component of Kalinzu chimpanzees.

### Isotopic variations among primate species

The comparison of hair isotope ratios of five primate species in Kalinzu indicated the lower δ^13^C_bulk_ and the lowest δ^15^N_bulk_ values in chimpanzees (Figure 1b, Tables 3 and 4). This result suggests little dietary contribution of animal proteins, which typically show higher δ^13^C_bulk_ and δ^15^N_bulk_ values because of higher TPs, in chimpanzees’ diet. The CSIA results also support the relatively low contribution of animal proteins in Kalinzu chimpanzees (Figure 4; Table 6). Previous studies have reported the successful reconstruction of meat consumption using bulk isotope analysis of bone collagen in Taï chimpanzees (Fahy et al., 2013), although not from the bulk hairs (Oelze et al., 2020). Our results, on the contrary, demonstrated that in Kalinzu chimpanzees, both bulk and CSIA analysis of hair were effectively employed to assess the relatively lower dietary contribution of animal proteins. Nevertheless, it should be acknowledged that research on meat-eating behaviors has been scarce, and previous systematic studies have focused on the extent of ant-eating behaviors (Hashimoto et al., 2000, 2015; Koops et al., 2015; Tashiro et al., 1999; Hashimoto et al., 2001; Ihobe, 2001; Shirasawa et al., 2024).

A L’Hoest’s monkey showed the highest δ^13^C_bulk_ and δ^15^N_bulk_ values among the primate species in Kalinzu (Figure 1b; Table 3). This is consistent with the visually observed feeding behaviors of L’Hoest’s monkeys, whose daily activities are mostly spent in the lower strata of the forest, like understray, and a large proportion of their diet consists of insects (Tashiro et al., 2006). Considering that the δ^15^N_bulk_ values of plants from undertray are significantly high in this site (Table 2) and the fact that insects generally show higher isotope ratios than plants (e.g., Hyodo et al., 2010), the isotopic signatures in L’Hoest’s monkey were consistent with their behavioral traits. The TP of 2.4, a slight faunivory, calculated in L’Hoest’s monkey in Kalinzu (Table 5) supports this interpretation. Relatively higher δ^15^N_bulk_ values in blue monkeys and red-tailed monkeys compared with chimpanzees (Figure 1b; Table 3) agree with the higher contributions of insects in their diet (Struhsaker, 2017; Tashiro et al., 2006).

The TP 2.0 measured by CSIA accurately reflects that guerezas are obligate leaf-eaters. On the other hand, guerezas’ bulk isotope ratios are significantly higher than those of chimpanzees (Table 4) and are the second highest after L’Hoest’s monkeys (Figure 1). The larger intra-species variations of δ^13^C_bulk_ and δ^15^N_bulk_ values in guerezas are also notable (Figure 1b; Table 3). Considering their heavy reliance on a limited variety of plant leaves (Matsuda et al., 2020), unknown physiological mechanisms might affect guerezas’ relatively higher and variable bulk isotope ratios. The investigation into the dietary, ecological, and physiological determinants of stable isotope ratios in guerezas is an interesting theme for future isotopic studies.

Across the primate species in Kalinzu, hairs from infants showed significantly higher δ^15^N_bulk_ values (Table 4). Infants show higher δ^15^N_bulk_ values (2–3‰) during the breastfeeding period because of breast milk consumption (Fahy et al., 2014; Tsutaya and Yoneda, 2015), which caused the significantly higher δ^15^N_bulk_ values in infant primate individuals in Kalinzu.

### Intra-individual variations

Rapid, large fluctuations of stable isotope ratios (>1‰) within several days were observed in the ultra-fine time-series analysis of single hair employed in this study, which was not previously observed in the sequential analysis of non-human primate hairs (e.g., Oelze et al., 2016). In particular, δ^15^N_bulk_ values showed a fluctuation (1.4‰) over a period of only 6 days (Figure 4; Supplementary Table S5) that was comparable to the largest difference (1.5‰) between chimpanzee individuals analyzed in Kalinzu (Table 3). If the fine-scale values over a period of several weeks are combined, the variation in isotope ratios from a single individual becomes much smaller (Figure 3; Supplementary Table S6). Stable isotope ratios, therefore, provide stable results as an average indicator of diet over a relatively long period of time, such as months or years. However, it should be noted that such average results can completely mask large and rapid potential fluctuations in isotope ratio, as shown in this study. Similar issues have been reported in other tissues with incremental growth, such as shells (Burchell et al., 2013) and teeth (Tsutaya, 2020). As recent analyses with higher temporal resolution are increasingly applied, the temporal resolution of stable isotope analysis may come closer to the daily resolution of behavioral observations, revealing previously entirely unanticipated intra-individual variations in isotope ratios and nutrient intake.

When using behavioral observation, collecting dietary data continuously over several months for a single individual can be difficult, and ultra-fine time-series isotope analysis offers a new solution. In our dataset, 64 days (73.6%) of the 87-day period corresponding to the hairs analyzed were not accompanied by an observational record of Kobo (Supplementary Table S8), and the diet during this period was not observed. Our analysis revealed unexpected isotopically fluctuating diet in Kobo, even for the periods without behavioral observational data. Further investigations into the factors affecting intra-individual isotopic variation, such as the rate and extent to which body nitrogen pools reflect inputs, physiological factors that can deviate hair growth, and fine-scale time-series changes in baseline isotope ratios, enable more nuanced interpretations of the ultra-fine time-series data.

Stable isotope analysis of primate hair has traditionally focused on dietary differences between individuals, but by applying ultra-fine time-series analysis, we can add a detailed temporal dimension to this analysis. This enables the investigation of interspecific and intraspecific dietary niche competition and differentiation in relation to seasonality of food resources and social interactions among individuals. Such investigations are applicable to subjects that are well-suited for stable isotope analysis, such as unhabituated populations and museum specimens, where direct behavioral observation is difficult. Improving the precision and the throughput of sectioning will be important in the future to implement these applications more effectively. Because the length of the hair segments to be sampled in ultra-fine sectioning is less than one-fifth to one-tenth the length compared with the conventional analysis, even the same degree of uncertainty in the length of the hair segments results in a relatively larger difference in the assigned time window. Because of the limited precision of manually cutting hairs into segments, it is necessary to use a microtome or other precise machine to section the hairs to finely control the time window represented by the hair segment. Such a partial automation of sectioning would also improve the throughput and enable a large-scale analysis.

## Conclusions

Carbon and nitrogen stable isotope analyses were performed on the hair and food plants of primates in Kalinzu to profile the isotopic variations. The baseline δ^13^C values in Kalinzu were >1‰ lower than those predicted from the rainfall. The δ^13^C and δ^15^N values of chimpanzees were generally lower than those of other primate species, and CSIA showed their plant-based diet, indicating the relatively small contribution of animal foods in the protein intake in Kalinzu chimpanzees. Ultra-fine time-series analysis of an adult male chimpanzee’s hair revealed large, rapid, unexpected intra-individual variations. This study showed that by improving and carefully implementing stable isotopic methods, detailed data comparable to behavioral observations can be obtained in primate ecology.

## Acknowledgments

We thank Takeshi Furuichi, Hiroshi Ihobe, Ikki Matsuda, Hodaka Matsuo, Daisuke Shimizu, Hiroyuki Takemoto, Haruka Taniguchi, Yasuko Tashiro, and Nahoko Tokuyama for supporting sample collection. This study was supported in part by Grant-in-Aid for Scientific Research (KAKENHI: 15J00464 and 25257409) from the Japan Society for the Promotion of Science.

## Author contributions

Conceptualization: TT, NA, CH; Methodology: TT; Software: TT; Investigation: TT, NFI, YS, MY, NOO, NO, CH; Resources: NA, HK, CH; Writing–Original Draft: TT; Writing– Review and Editing: TT, NFI, HK, NOO, NO, CH; Visualization: TT; Funding acquisition: TT, CH.

## Notes

### Competing Interest Statement

The authors have declared no competing interest.

